# ToolsGenie 2.0: A Scalable and Extensible Multi-Agent System for Bioinformatics Automation

**DOI:** 10.64898/2026.01.06.697527

**Authors:** Youjia Ma, Bo-Wei Han, Minzhe Zhang, Yang Leng, Wenhao Gu, KC Shashidhar, Xiao Yang

## Abstract

The rapid expansion of biomedical data necessitates efficient bioinformatics tools, yet conventional workflows rely heavily on manual dependencies, hindering scalability and broader adoption. Building on the foundation of ToolsGenie 1.0, we introduce ToolsGenie 2.0, a multi-agent AI framework that automates bioinformatics analyses through natural language queries and file inputs. ToolsGenie 2.0 addresses the growing need for customizable analyses by offering extensibility, reproducibility, improved accuracy, and ease of use. It incorporates a ReAct-based architecture with dual-layer extensibility for sub-agents and specialized tools, along with dynamic Docker image selection for automated, sandboxed, and secure environment management. Rigorous benchmarking shows that ToolsGenie 2.0 achieves 68.6% accuracy on an in-house dataset and 48.3% accuracy on BixBench, demonstrating competitive performance across diverse evaluation settings. These innovations position ToolsGenie 2.0 as a versatile platform for broadening access to bioinformatics in both research and clinical contexts. ToolsGenie 2.0 is available on the PromptBio platform (<u>platform.promptbio.ai</u>).

## Introduction

The biomedical field has witnessed an exponential growth in data generation, driven by advancements in high-throughput technologies such as next-generation sequencing, proteomics, and metabolomics^1,2^. This surge in biological data necessitates sophisticated bioinformatics analysis to extract meaningful insights, enabling breakthroughs in disease understanding, drug discovery, and personalized medicine. However, traditional bioinformatics workflows often rely heavily on manual intervention, requiring domain experts to select appropriate tools, configure parameters, manage computational environments, and troubleshoot errors^3^. This reliance on human expertise not only introduces scalability and reproducibility bottlenecks but also exacerbates challenges for non-specialists, leading to inefficiencies, higher error rates, and barriers to widespread adoption in resource-limited settings.

To bridge the gap, several LLM-powered and agentic systems have emerged to automate parts of the bioinformatics stack, marking incremental advancements in tackling the field’s complexities. **AutoBA** (Automated Bioinformatics Analysis)^4^ and **BioMaster**^5^ represent early large language model (LLM)–based approaches to bioinformatics data analysis. In both systems, users provide datasets and task requirements upfront, after which the agent autonomously generates and executes code to complete the analysis. However, despite their strengths in workflow automation, neither system supports multi-round conversational interaction or real-time user engagement during analysis, making them incompatible with tasks that require interactive, dialogue-driven decision-making. **Biomni**^6^ adopts a ReAct-style agent architecture^7^ to function as a general-purpose biomedical AI, integrating a wide range of specialized tools and databases to execute diverse research tasks through natural language interaction. **STELLA**^8^ extends this paradigm by emphasizing self-evolution, enabling the agent to autonomously discover, evaluate, and incorporate new tools over time, thereby adapting to novel problem domains and expanding its capabilities dynamically. Beyond general-purpose bioinformatics agents, a complementary class of domain- or task-specialized code-executing agents has emerged. While not designed to handle arbitrary workflows, these systems can autonomously generate and execute analysis code within well-defined domains. For example, **CellAgent**^9^ enables end-to-end single-cell RNA-seq analysis via an LLM-driven multi-agent framework guided by natural language instructions, whereas **DrBioRight**^10^ targets large-scale cancer functional proteomics by integrating natural language querying, automatic code generation, and executable workflows for multi-omics datasets such as TCGA and CCLE. Collectively, these systems occupy an intermediate position between fully general-purpose agent frameworks and narrowly scoped analytical tools.

While these agents could substantially reduce manual effort, they often fall short in achieving operational flexibility and reliability. For example, they frequently face constraints in compatibility with expansive tool libraries and rely on rigid, predefined setups that do not accommodate the dynamic needs of bioinformatics analyses, where new software integrations can conflict with existing configurations^11^. Additionally, multi-agent designs enhance modularity but introduce heightened system complexity, rendering memory and state management fragile during prolonged interactions and increasing the risk of errors^6^. Similarly, while the ReAct architecture promotes more humanistic, conversational interactions, it often generates verbose contexts that can obscure user intent and hinder accurate analysis completion^7^. Furthermore, inadequate validation mechanisms for input, output, and intermediate files further exacerbate these issues, leading to execution failures or unreliable results, particularly when handling heterogeneous file formats and data structures or when the data size is large. Moreover, these agents typically fall short of industrial-grade standards, particularly in security and data integrity, as generated code is often executed within the same environment as the reasoning agent, alongside limited cost management, resource control, and reproducibility guarantees—factors that hinder scalable, production-level deployment.

We introduce **ToolsGenie 2.0**, an AI agent that autonomously performs bioinformatics analyses using user queries and input data files. Building on **ToolsGenie 1.0**^13^, its key objectives are to: (1) enable rapid prototyping from idea to production, (2) support customized analysis needs, and (3) improve analytical coverage, accuracy, and reproducibility. To achieve these goals and overcome limitations in existing tools, ToolsGenie 2.0 incorporates several innovations: (1) automated environment selection using Docker-based sandboxes for seamless execution across diverse infrastructures; (2) an extensive library of specialized bioinformatics tools for flexible and extensible integration; (3) multi-branch memory management to maintain a concise, stable state for the main agent, preventing overload and boosting decision-making; and (4) systematic file validation for inputs, outputs, and intermediates to ensure correctness and higher success rates. These features reduce human dependency, enhance reliability, and make ToolsGenie 2.0 a versatile tool for diverse biomedical research applications.

### Core Contributions

#### Extensible ReAct multi-agent framework

ToolsGenie 2.0 implements a modular ReAct-style multi-agent architecture with dual-layer extensibility (sub-agents and tools)^7,14^, enabling rapid addition of new analytical capabilities while keeping planning and tool invocation interpretable and debuggable.

#### Industrial-grade execution with Docker sandboxes and cost-aware memory

The system automatically selects Docker-based sandbox environments for reliable, portable execution and employs multi-branch memory management that preserves task context concisely to reduce token usage and operational cost, while also storing executed code to support reproducible, iterative analyses.

#### Evaluation on public and in-house benchmarks

We validate ToolsGenie 2.0 on the BixBench dataset^15^ and an in-house dataset comprising a range of bioinformatics analysis tasks, showing competitive performance across both public and real-world cases.

#### Ready-to-use deployment

ToolsGenie 2.0 is designed as a cloud-native agent and is deployed on the PromptBio platform (platform.promptbio.ai)^13^, enabling users to run complex, reproducible analyses directly from natural-language queries and file inputs without any need for a local setup.

## Results

### ToolsGenie 2.0 Framework

The overall architecture of **ToolsGenie 2.0** is illustrated in Figure 1. Users submit analysis requirements and input data, which are processed by a central **Supervisor Agent** built on a ReAct-style architecture that tightly integrates reasoning and action. The Supervisor Agent generates a structured analysis plan and dynamically orchestrates downstream task execution by selectively invoking specialized sub-agents—namely the **ExecutorAgent, KnowledgeAgent**, and **ReferenceAgent**—to progressively complete the workflow (see Methods). This ReAct-driven control loop enables the Supervisor Agent to iteratively reason over intermediate results, refine decisions, and invoke appropriate tools or sub-agents as the analysis progresses.

**Figure 1.**
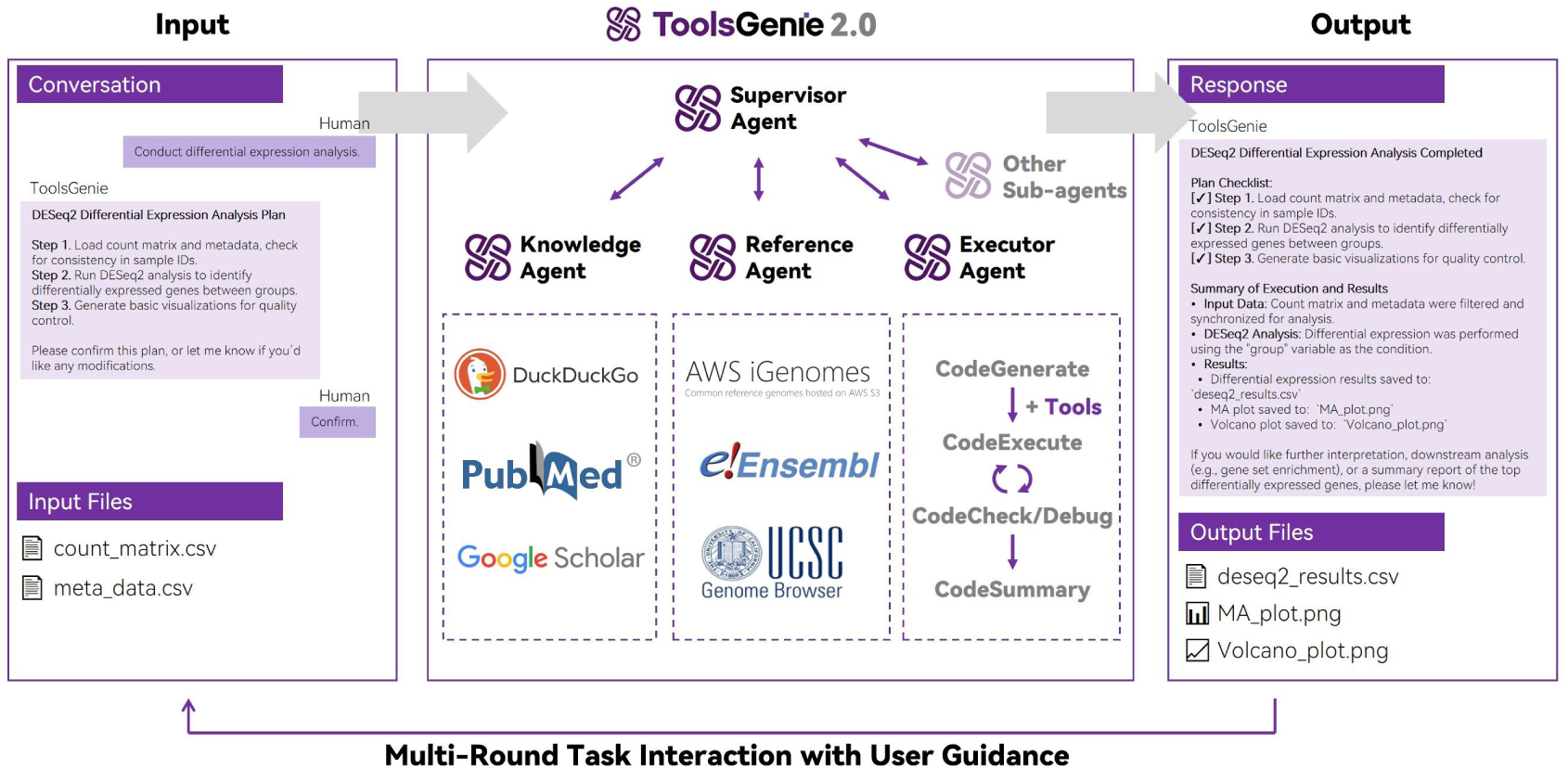
Overview of ToolsGenie 2.0.

Because most bioinformatics workflows require custom data processing code, the ExecutorAgent plays a central role in ToolsGenie 2.0. It autonomously performs task-specific operations through an iterative cycle of code generation, execution, and debugging. A major challenge in bioinformatics analysis lies in software and environment configuration, where static or preconfigured setups frequently fail due to dependency conflicts—particularly when required tools are absent from the base installation. To address this, ToolsGenie 2.0 decomposes tasks during the planning phase and, upon ExecutorAgent invocation, dynamically selects and deploys appropriate Docker images from container repositories such as Docker Hub, Quay.io, and Wave. This strategy avoids installation bottlenecks while leveraging container sandboxing to ensure execution isolation, enhanced data security, granular permission control, and protection from host system vulnerabilities.

To support scalability and extensibility, the ExecutorAgent adopts a standardized execution interface based on **LangGraph**, which simplifies tool management and invocation during analysis. This design enables seamless integration of large, specialized tool ecosystems from platforms such as **Biomni** and **STELLA**, allowing ToolsGenie 2.0 to incorporate new analytical tools and workflows as bioinformatics methodologies evolve. Together, these execution and extensibility mechanisms ensure robust handling of diverse, multi-stage analyses while maintaining adaptability to emerging computational needs.

Beyond execution, ToolsGenie 2.0 incorporates knowledge- and resource-aware agents to enrich analytical context and facilitate standardized data access. The KnowledgeAgent enables internet-based queries, including web content retrieval and scholarly literature searches, allowing the system to incorporate external contextual information during analysis. Complementarily, the ReferenceAgent retrieves essential public resources—such as reference genomes or pre-indexed files for alignment tools (e.g., BWA and STAR)—from repositories including AWS iGenomes. Together, these agents augment execution with domain knowledge and standardized resources, enabling more informed and reproducible bioinformatics workflows.

A key enabling component of the ToolsGenie 2.0 framework is its **contextual memory management system**, which regulates information flow across agents and supports multi-round interactions. The Supervisor Agent maintains a dedicated memory branch for task-level context, including user interactions, conversation history, high-level plan updates, and sub-agent invocations. Task-specific artifacts—such as file metadata, intermediate results, executed code, and configuration parameters—are stored separately in a shared **LangGraph state**. Sub-agents access only the information required for their operations and return essential outputs to update the shared state for subsequent steps. This organization prevents context dilution, reduces latency in complex workflows, and improves scalability, while enabling prior analytical steps to be reliably reused or re-executed by preserving executed code as a first-class artifact within the shared state—either verbatim or with new input data—thereby supporting reproducible and iterative analysis.

Importantly, this memory design enables **multi-turn, human-in-the-loop analysis** by preserving task context across successive user interactions (Figure 1). Outputs from one round can be reviewed, refined, or corrected by the user and reused as inputs in subsequent rounds, allowing iterative analysis without loss of contextual coherence. By explicitly segregating task-level and execution-level context and regulating information flow between them, ToolsGenie 2.0 ensures that complex, multi-round bioinformatics analyses remain efficient, accurate, and robust.

The core supervisor agent, based on a ReAct architecture, processes user conversations and input files to create a high-level analysis plan. This plan is executed by selectively invoking specialized core sub-agents, including the **ExecutorAgent**, which autonomously handles task-specific code generation, execution, and debugging, as well as the **KnowledgeAgent** for web and literature searches and the **ReferenceAgent** for retrieving standardized public resources. The framework offers extensibility through additional sub-agents for domain-specific or utility tasks and integration of external tools, facilitating adaptation to various bioinformatics scenarios. Outputs from each round are reused as inputs in subsequent rounds, enabling multi-round, human-in-the-loop iterative analysis through user review, guidance, and correction.

### Docker Image Selection Enhances Accuracy and Efficiency in Bioinformatics Tasks

To demonstrate how ToolsGenie 2.0’s dynamic Docker image selection mechanism improves task accuracy and efficiency, we evaluated it on 10 representative common bioinformatics tasks that require third-party software or libraries (including R packages, Python packages, and shell command-line tools). These tasks contain the four most frequently used steps in genomics analysis and the four most commonly applied steps in transcriptomics analysis, alongside a typical sequence processing task and a machine learning task—selected specifically to reflect real-world scenarios where third-party software and libraries necessitate manual installation and configuration.

For each task, we compared ToolsGenie 2.0 with its Docker image selection enabled—dynamically retrieving optimized images from repositories including Docker Hub, Quay.io, and Wave containers—against baseline configurations that relied on generic base images (“r-base:4.5.1” for R-dependent tasks, and “condaforge/miniforge3:25.11.0-0” for Python- and Shell-dependent tasks). In the baselines, the ExecutorAgent attempted to automatically install required tools or libraries during execution within these generic base images. All evaluations were conducted on an Ubuntu server equipped with Intel Xeon Gold 6238R CPUs (112 logical cores, 2.20 GHz) and 629 GB of RAM. Resource availability remained sufficient throughout the evaluations. Given the relative simplicity of the tasks, we employed the GPT-4.1 model for the analysis. Each task was repeated three times to account for variability. Accuracy was defined as successful task completion with validated correct outputs, where executions finished without errors and produced expected results for the corresponding bioinformatics tool. Efficiency was quantified by average execution time (in seconds). The selected tasks are detailed in Table 1.

**Table 1.** Ten Representative High-Frequency Bioinformatics Tasks Requiring Third-Party Software/Libraries.

| Task ID | Task Description | Dependency Type | Required Tools/Libraries |
| --- | --- | --- | --- |
| Q1 | Align sequencing reads from FASTQ files using BWA. | Shell | BWA |
| Q2 | Mark duplicates in BAM files using Picard. | Shell | Picard |
| Q3 | Perform germline variant calling using GATK. | Shell | GATK |
| Q4 | Generate a TBI index file for the VCF.GZ file. | Shell | Tabix |
| Q5 | Conduct differential expression analysis from count matrices using DESeq2. | R | DESeq2 |
| Q6 | Estimate immune cell proportions in bulk RNA-Seq data using xCell. | R | xCell |
| Q7 | Perform gene set enrichment analysis (GSEA) from expression data using the KEGG pathway database with clusterProfiler. | R | clusterProfiler |
| Q8 | Perform clustering and generate UMAP plots colored by cluster identities using Seurat. | R | Seurat |
| Q9 | Translate an mRNA sequence into a protein sequence from the first start codon using BioPython. | Python | Biopython |
| Q10 | Train a random forest regressor to predict patient age from DNA methylation levels (beta values) and select the top 20 features using scikit-learn. | Python | scikit-learn |

Comparative results demonstrate that the Docker image selection mechanism substantially enhances both success rates and efficiency. Across the 10 tasks, ToolsGenie 2.0 achieved an average success rate of 83.3%, compared to 56.7% for the baseline (Figure 2A). Notably, one exception was observed in Task Q5, where the accuracy achieved with Docker image selection was lower than that of the baseline. Upon manual inspection, this discrepancy was traced to an occasional code-generation error by the LLM, which produced an incorrect DESeq2^16^ analysis script in that specific run, rather than to the Docker-based execution environment itself. Crucially, this variability is due to the inherent non-determinism of LLM outputs, not to the reproducibility of the workflow itself: in the subsequent three repeated executions of Task Q5 under the same Docker configuration, the error did not reappear and correct results were obtained. While this isolated discrepancy affected task accuracy in a single case, efficiency gains were particularly pronounced: the average execution time with Docker selection was 257.3 seconds per task, versus 913.0 seconds for the baselines, with the reduction primarily attributable to the elimination of installation delays (Figure 2B). This underscores the inherent challenges of software installation in bioinformatics analyses—a step that is notoriously demanding due to complex dependencies, compilation requirements, and potential conflicts with existing environments. Certain tools, particularly R packages, are difficult to install automatically; they often involve intricate dependencies, extensive compilation times, and incompatibilities that can lead to failures. For instance, popular packages such as xCell^17^, clusterProfiler^18^, and Seurat^19^ failed to install successfully in the baseline’s clean “r-base:4.5.1” image, despite repeated attempts. These failures were exacerbated in environments pre-populated with other software, where conflicts with existing libraries or system configurations further compounded the issues, leading to incomplete executions or crashes. By contrast, ToolsGenie 2.0’s Docker image selection bypasses these pitfalls by dynamically sourcing pre-configured, optimized images that already include the necessary tools and resolved dependencies, thereby ensuring more robust and streamlined workflows.

**Figure 2.**
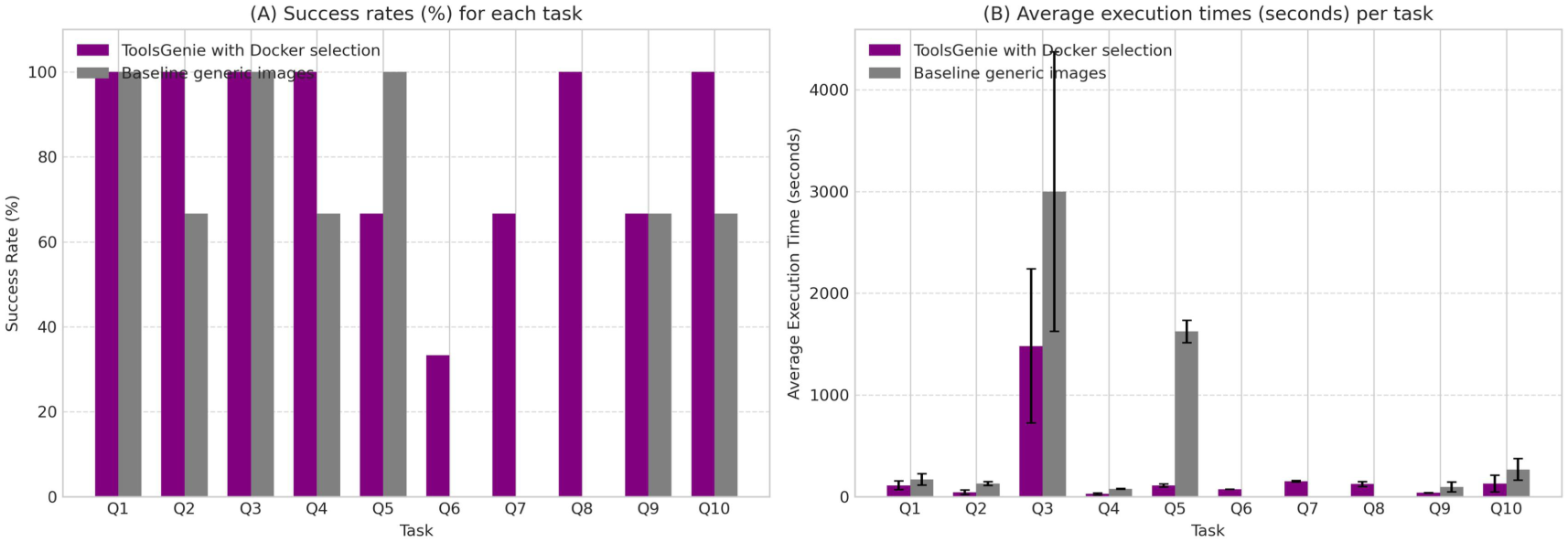
Comparative Performance of Docker Image Selection Mechanism. (A) Success rates (%) for each task, with bars representing ToolsGenie 2.0 with Docker selection (purple) versus baseline generic images (gray; aggregated across R-base, Python-base, and Ubuntu-base where applicable). Higher bars indicate better reliability, with Docker selection outperforming baselines by mitigating installation failures. (B) Average execution times (seconds) per task, showing reduced processing durations for Docker selection (purple) compared to baselines (gray), attributed to avoided dependency resolution overhead. Execution time averaged over 3 runs per task, with only the times of successfully executed tasks being incorporated into the calculation; error bars denote standard deviation. The figure was generated by ToolsGenie 2.0 based on the figure legend and corresponding data (Supplementary Note 3).

### Sub-Agents and Specialized Tools Expand ToolsGenie 2.0’s Capabilities Across General and Domain-Specific Bioinformatics Tasks

To further demonstrate how ToolsGenie 2.0 scales beyond general-purpose workflows and effectively handles domain-specific analyses, we evaluated the contributions of extended sub-agents and tools.

First, we assessed the impact of the ReferenceAgent on tasks requiring reference genomes or annotation resources—elements that are crucial in bioinformatics but often overlooked or inadequately supported by existing bioinformatics agents. We selected a representative task: RNA-seq data alignment, which requires the accurate retrieval or construction of large reference files (e.g., FASTA sequences, GTF annotations, and pre-built indexes). For analyses involving mouse samples, enabling the ReferenceAgent allowed ToolsGenie 2.0 to autonomously fetch the correct reference data from AWS iGenomes and seamlessly integrate it into downstream workflows (Supplementary Note 1A). In contrast, when the ReferenceAgent was disabled, ToolsGenie 2.0 resorted to downloading the necessary reference files using the KnowledgeAgent or ExecutorAgent, followed by building STAR^20^ indices directly from the FASTA file (Supplementary Note 1B). This not only substantially prolonged the analysis time and cost (as building an index is typically a time-consuming and memory-intensive process) but also elevated the risk of errors: download URLs provided by the LLM were often incorrect (particularly for less commonly used references), and web searches could encounter reliability issues. Therefore, in specialized fields like bioinformatics, extending the system with dedicated sub-agents for tasks such as handling reference genomes, accessing specific databases, managing complex annotations, processing high-throughput data, and performing specialized analyses can significantly enhance efficiency, reduce computational overhead, minimize error rates, improve reproducibility, and ensure industrial-level reliability by enabling reliable, autonomous handling of domain-specific resources.

Next, to evaluate ToolsGenie 2.0’s capacity for addressing specialized tasks beyond the capabilities of general-purpose LLM-based code synthesis, we examined its performance when augmented with externally curated toolsets. Using the Biomni toolkit^7^ as a case study, we tested ToolsGenie 2.0’s ability to predict ADMET properties for the compound CC(C)CC1=CC=C(C=C1)C(C)C(=O)O. Predicting ADMET properties is a challenging task requiring specialized computational models, extensive chemical databases, and domain-specific algorithms to accurately assess Absorption, Distribution, Metabolism, Excretion, and Toxicity—factors essential for drug discovery but often inaccessible through generic tools. Without access to domain-specific tools, the baseline system struggled to complete the analysis, including erroneously invoking non-command-line tools (Supplementary Note 2C), attempting direct web searches for answers (Supplementary Note 2D), and even when leveraging RDKit software to obtain results, producing outputs with inconsistent formats and content that may omit essential information (Supplementary Notes 2E and F)—rendering it unsuitable for real-world production environments. In contrast, after integrating the Biomni toolkit module, ToolsGenie 2.0 successfully and consistently completed the task, maintaining output consistency even across different models (e.g., GPT-4.1 and Claude-4-Sonnet), thereby ensuring correct and reproducible outputs that can be reliably applied in production settings (Supplementary Notes 2A and B). These findings illustrate that ToolsGenie 2.0’s tool integration facilitates sophisticated, domain-specific workflows that would otherwise be infeasible for a general-purpose agent and enables continuous extension of its tool ecosystem as new tools become available.

Taken together, these evaluations demonstrate that the combined design of sub-agents and specialized tool integrations fundamentally enhances ToolsGenie 2.0’s breadth and depth.

### Performance Evaluation of ToolsGenie 2.0

To benchmark ToolsGenie 2.0’s effectiveness, we compared it against Biomni, a state-of-the-art bioinformatics agent, using both an in-house curated dataset and the publicly available BixBench dataset (version 1.5) )^15^. All evaluations were conducted with the Claude-4-Sonnet model to ensure consistency.

Performance was quantified by accuracy, defined as the proportion of tasks yielding outputs that aligned with ground-truth or expected results. Our in-house curated dataset comprised 140 diverse bioinformatics questions across various types and scenarios (Supplementary Table 1), each including a query, input data, and reference answers generated by human bioinformaticians. Outputs from both agents were independently assessed by three bioinformatics experts. On this dataset, ToolsGenie 2.0 achieved an accuracy of 68.6%, outperforming Biomni’s 60.0% (Figure 3A). This improvement stemmed primarily from ToolsGenie 2.0’s Docker image selection mechanism, which enabled the successful execution of numerous bioinformatics tools that Biomni failed to install or utilize (Supplementary Table 1). Notably, the average cost per task on this dataset was comparable between the two agents ($0.51 for ToolsGenie 2.0 versus $0.52 for Biomni; Figure 3B), indicating that the accuracy gains were achieved without additional cost overhead.

**Figure 3.**
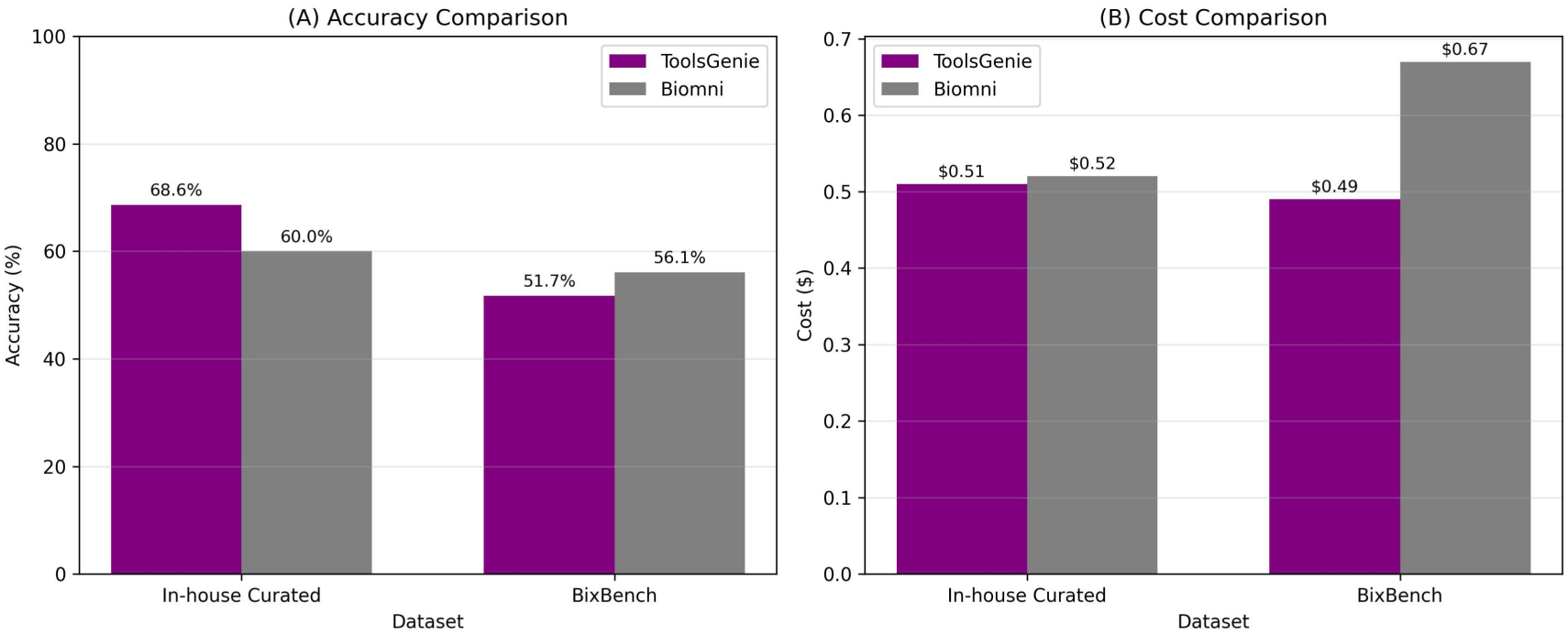
Performance comparison between ToolsGenie 2.0 and Biomni.

On the BixBench dataset, which comprises 205 bioinformatics data analysis questions formatted as multiple-choice questions, ToolsGenie 2.0 achieved an accuracy of 51.7% (106/205), slightly below Biomni’s 56.1% (115/205; Figure 3A and Supplementary Table 2). To identify the primary cause of this performance difference, we conducted a post-hoc analysis of the 33 cases in which Biomni produced correct answers while ToolsGenie 2.0 did not. Among these, 15 cases were traced to only two datasets characterized by exceptionally large numbers of input files. Specifically, seven questions were associated with a dataset containing 88 Excel files, while eight questions originated from a dataset consisting of 11 compressed archives that, once decompressed, yielded 3842 files spanning 20 distinct file formats. In these scenarios, ToolsGenie 2.0’s ExecutorAgent encountered limitations during file inspection and summarization. The LLM-generated scripts produced by the ExecutorAgent, which were used to enumerate file contents and extract structural information, generated outputs that exceeded predefined length constraints, resulting in partial truncation of file-level details. In addition, when summarizing large collections of heterogeneous files, the LLM-based aggregation step occasionally omitted critical information. As a result, incomplete data descriptions were propagated to downstream agents, leading to suboptimal analysis plans and incorrect answers. These errors therefore arose primarily from information loss during large-scale file inspection and inter-agent information transfer, rather than from deficiencies in tool execution or analytical reasoning. By comparison, Biomni-A1’s single-agent approach preserves information coherence within a single interaction, which may explain why it correctly answered these cases.

Beyond accuracy, we also evaluated average inference cost per task. From a cost perspective, ToolsGenie 2.0 consistently incurred lower average inference costs on the BixBench dataset ($0.49 per task) compared to Biomni ($0.67 per task; Figure 3B). This observation suggests that, despite reduced accuracy under extreme input-scale conditions, ToolsGenie 2.0 maintains favorable cost characteristics at the system level.

Bar plots comparing the performance of ToolsGenie 2.0 and Biomni across two datasets: an in-house curated dataset and the publicly available BixBench dataset. Accuracy, defined as the correctness of outputs, is shown in (A), and average cost is shown in (B). For each dataset, two bars represent the two agents (purple: ToolsGenie 2.0; gray: Biomni). The figure was generated by ToolsGenie 2.0 based on the figure legend and corresponding data (Supplementary Note 4).

Moreover, we attempted to evaluate STELLA on the BixBench dataset. During initial runs of seven selected questions, two tasks (bix-10-q1 and bix-10-q5) each required 7.59 and 7.81 hours to complete, preventing us from processing the full dataset. While STELLA performed well on these seven tasks—correctly answering three questions, compared to one correct answer each for ToolsGenie 2.0 and Biomni—the average runtime (2.30 hours per task) and token consumption (mean input 308,323 tokens, mean output 40,118 tokens) were substantially higher than ToolsGenie 2.0 on the same questions (average runtime 5.0 minutes, mean input 98,623 tokens, mean output 14,873 tokens). Given that LLM selection is a key determinant of agent performance, and STELLA employs a mixture of three models (Gemini-2.5-Pro, Claude-Sonnet-4, and Grok-4), a fair comparison of agent architectures is not feasible. Consequently, STELLA was not included in the full-scale benchmark.

Overall, these results highlight that ToolsGenie 2.0 achieves performance comparable to that of state-of-the-art agents across a range of benchmarks, particularly in domains demanding robust tool integration and reliable execution of intricate workflows.

## Discussion

In this work, we present ToolsGenie 2.0, a multi-agent framework designed to tackle the complexities of bioinformatics analysis. This system enables rapid prototyping of solutions with broad analytical coverage, accuracy, and reproducibility through modular extensibility. Our experimental evaluations demonstrate that ToolsGenie 2.0 not only performs on par with existing frameworks like Biomni but also offers scalable, engineering-oriented features for real-world deployment.

Although LLMs have demonstrated increasingly powerful coding abilities, persistent pain points remain, particularly in environment setup in complex domains^11,21^. ToolsGenie 2.0 mitigates these issues through a dynamic Docker image selection mechanism, which not only enhances efficiency by bypassing manual installations but also promotes reproducibility. However, the mechanism’s effectiveness relies on the coverage and quality of community-maintained Docker repositories. Incomplete or outdated images could introduce vulnerabilities, underscoring the need for ongoing community contributions to maintain a comprehensive library of pre-configured bioinformatics environments. Therefore, developing more advanced methods for automated Docker image configuration would be a valuable and necessary advancement to further reduce environment-induced task failures.

A core strength of ToolsGenie 2.0 lies in its modular architecture, which enables seamless extension via sub-agents and specialized tools, allowing adaptation to emerging research needs and user preferences. This dual-layer extensibility positions ToolsGenie 2.0 as a versatile platform for both routine and advanced analyses. Looking ahead, we envision integrating domain-specific sub-agents for fields like single-cell transcriptomics or protein structure prediction, embedding standardized pipelines and best-practice parameters to boost accuracy, standardization, and reproducibility. Utility sub-agents, such as a ReportGenerationAgent for structured outputs with charts and explanations or a CodeSummaryAgent for natural language code abstracts, could further enhance user-friendliness. Moreover, ToolsGenie 2.0 could complement established best-practice workflows like nf-core pipelines^22^ or Snakemake Workflows^23^ by transforming these pipelines into sub-agents or tools. Thus, bridging exploratory analysis with scalable production, enabling a hybrid paradigm where natural language drives standardized, reproducible pipelines.

Nonetheless, several limitations remain. In practical bioinformatics settings, ToolsGenie 2.0 demonstrates strong potential but faces constraints with large-scale data and input variability. For massive datasets—characterized by both high file counts and large individual file sizes—the sheer volume of file descriptions can consume substantial context tokens, leading to context overflow and diluted attention in LLMs. This issue is particularly detrimental in multi-agent architectures, where information must be passed between agents, potentially degrading task completion quality. In the BixBench dataset, certain problems involve hundreds of files in diverse formats, and the token overload during inter-agent transmission resulted in information loss, contributing significantly to ToolsGenie 2.0’s suboptimal performance on this benchmark. Additionally, LLM-generated code introduces additional challenges in both efficiency and stability. In our experiments, such code sometimes failed to fully leverage available computational resources, resulting in prolonged runtimes and reduced throughput. Occasional non-deterministic errors—where code produced transient mistakes in specific runs—can also affect workflow robustness, even when the underlying execution environment is properly configured. These characteristics highlight the importance of evaluating LLM-driven workflows with multidimensional metrics beyond average success rates, including consistency, reproducibility, and resource utilization. Current benchmarks, such as BixBench, while valuable, often overlook real-world complexities such as noisy data or iterative user refinements^24^. We advocate developing more ecologically valid benchmarks that simulate authentic research scenarios, incorporating multi-turn dialogues and heterogeneous inputs to better assess agent utility.

ToolsGenie 2.0 holds transformative potential in democratizing bioinformatics, lowering barriers for non-experts by enabling wet-lab researchers or clinicians to conduct complex analyses via natural language, shortening the path from data to insights in a flexible and reproducible manner. This is particularly impactful in resource-constrained settings, such as small laboratories or institutions, where computational infrastructure and expertise are limited, thereby fostering scientific equity. We foresee AI-driven, natural language interfaces becoming the norm in bioinformatics, fundamentally altering workflows and enabling autonomous scientific discovery systems. ToolsGenie 2.0 represents a pivotal step toward this vision, empowering researchers to focus on innovation rather than technical hurdles.

## Methods

### Multi-Agent Framework

ToolsGenie 2.0 is a multi-agent system developed using LangGraph. The architecture comprises a central Supervisor Agent based on the ReAct architecture^7^, along with several core sub-agents: the KnowledgeAgent, ReferenceAgent, and ExecutorAgent. This modular design enables autonomous orchestration of bioinformatics workflows, allowing the system to decompose complex user queries into actionable steps while maintaining efficiency and adaptability.

The Supervisor Agent serves as the orchestrator, leveraging user-provided analysis requirements and pre-summarized file metadata to generate and execute a high-level plan.

Inputs to the Supervisor include historical conversation context and uploaded files. For files in tabular formats (e.g., CSV, TSV, TXT) or bioinformatics-specific formats (e.g., FASTQ, FASTA, BAM, BED), ToolsGenie 2.0 employs predefined parsing logic to rapidly extract essential summarization without loading the entire content into memory. The Supervisor Agent first assesses user intent to determine the response strategy: for simple queries—such as greetings or basic operations like “computing the reverse complementary sequence of ATCCGGCCAAGT”—it directly generates a response. For more complex tasks, it follows the ReAct paradigm, iteratively reasoning about the plan, invoking appropriate sub-agents as actions, and observing their outputs to refine subsequent steps. Upon completion of all planned actions, the Supervisor Agent synthesizes the results into a coherent summary and delivers it to the user, ensuring end-to-end traceability.

The KnowledgeAgent addresses the inherent limitations of LLM parametric knowledge by providing dynamic access to external information sources through querying web-based resources. It utilizes search APIs to retrieve relevant webpages, parses their content, and summarizes key insights tailored to the user’s query. Thereby, enrich the analysis plan with up-to-date, domain-specific information and mitigate gaps in the LLM’s internal knowledge.

The ReferenceAgent addresses a common bottleneck in bioinformatics pipelines: the acquisition of large reference datasets, which often involve time-consuming downloads and indexing processes for tools like BWA or STAR. This agent searches local caches or public repositories such as AWS iGenomes for pre-built resources (e.g., reference genomes, annotation files, or pre-indexed alignment databases). Based on the user’s query and task requirements, it identifies and retrieves the most suitable references, returning their metadata and access paths to the Supervisor Agent for integration into subsequent steps, thus accelerating workflows and reducing computational overhead.

The ExecutorAgent is responsible for translating high-level analytical tasks into executable code and managing their execution through an iterative loop of code generation, execution, and debugging. It aggregates necessary file metadata, user specifications, and contextual information from the Supervisor Agent to prompt the LLM for generating executable code snippets in languages like Python, R, or Shell. The code is then run in a controlled environment, with standard output (stdout) and error (stderr) captured and fed back to the LLM for result verification and debugging in case of failures. The code is then run in a controlled environment, with standard output (stdout) and error streams (stderr) captured and fed back to the LLM for result verification, as well as for debugging in the event of failures. Task completion is validated through rigorous file checks on outputs and intermediates, including confirmation of file existence, non-empty content, and correct format. Upon successful validation, the agent provides a concise summary of the results and shares output file paths with the Supervisor Agent. To handle potential failures gracefully, a retry limit is enforced; if exceeded, the agent reports the issue back to the Supervisor Agent, prompting a strategy update such as task decomposition or alternative tool selection. This self-correcting mechanism enhances overall reliability and minimizes disruptions in multi-step analyses.

Notably, the Supervisor Agent operates with no direct access to modify user data, processing only metadata, task specifications, and summarized results. The ExecutorAgent operates with time-limited read-only access to specific input files and write-only access to designated output locations, ensuring that analytical code cannot access unauthorized files or modify original data. Collectively, these designs establish a secure execution boundary that is essential for handling sensitive biomedical data.

### Contextual and Execution-Aware Memory Management in ToolsGenie 2.0

ToolsGenie 2.0 implements a multi-branch memory management system to efficiently handle contextual information across its multi-agent architecture, preventing information overload and ensuring streamlined decision-making.

At the core, the Supervisor Agent maintains a dedicated memory branch for all user interactions, capturing conversation history and high-level plan updates in a concise format. In parallel, task-specific information—such as file metadata, intermediate results, executed code, and configuration parameters—is stored separately in the LangGraph shared state. During execution, sub-agents receive targeted guidance from the Supervisor Agent, including the current task description and relevant context from the user memory. They then pull necessary details directly from the LangGraph state to perform their operations. Upon completion, sub-agents return only essential outputs to the Supervisor Agent, while updating the shared state with any new artifacts for potential future reference. Notably, executed code and associated parameters are preserved in the shared state as reusable execution records, enabling subsequent re-execution or controlled reuse in later analysis rounds. This selective feedback mechanism ensures that the Supervisor Agent remains agnostic to the intricate internal workings of sub-agents, concentrating solely on integrating results, monitoring progress, and adjusting the overall plan as needed.

By decoupling user-level context from granular task data and enforcing minimal information flow between branches, ToolsGenie 2.0’s memory management enhances scalability, reduces latency in long-running analyses, and improves robustness against context dilution, while enabling prior analytical steps to be reliably reused or re-executed to support reproducible, multi-round bioinformatics workflows—ultimately leading to more accurate and efficient bioinformatics outcomes.

### Extensibility of Sub-Agents and Tools in ToolsGenie 2.0

ToolsGenie 2.0 is designed with a highly modular architecture that supports straightforward extension at two distinct levels: the addition of new sub-agents and the integration of external tools for use by the ExecutorAgent. This design allows users or developers to expand system capabilities based on specific needs without altering the core architecture (Figure 1).

At the sub-agent level, ToolsGenie 2.0 supports dynamic registration of new specialized agents using LangGraph’s standard tool-calling interface, enabling seamless integration into the Supervisor Agent’s invocation workflow. Domain-specific sub-agents and utility-oriented sub-agents can be added to incorporate standardized pipelines, improve accuracy, reproducibility, and enhance user experience.

At the tools level, the ExecutorAgent enables extension by directly generating and executing code to invoke external tools, rather than relying on LangGraph’s standard interface. It supports integration of extensive specialized toolsets from platforms like Biomni and STELLA through simple configuration of tool registration files, allowing broader task coverage during analysis.

This dual-layer mechanism ensures ToolsGenie 2.0’s broad compatibility with external tools, making it adaptable to evolving bioinformatics requirements.

### ToolsGenie 2.0’s Docker Image Selection Mechanism

To minimize software installation efforts and ensure compatibility in heterogeneous environments, ToolsGenie 2.0 incorporates a dynamic Docker image selection mechanism^25^. This mechanism enables ToolsGenie 2.0 to flexibly leverage a broad range of general-purpose bioinformatics tools and libraries, supporting the construction of custom workflows for diverse and previously unseen analytical requirements. During the code generation phase, the ExecutorAgent synchronously identifies required third-party software and libraries. At execution time, it automatically searches local caches and public repositories—such as Docker Hub, Quay.io, and Wave containers, which collectively encompass 115,258 indexed Docker images as of December 2025—for the image that most closely matches these dependencies, prioritizing minimalistic images with pre-installed tools to avoid conflicts. Complementing this, the Supervisor Agent optimizes the granularity of the generated plan by decomposing complex tasks into smaller, focused steps, ensuring that each invocation of the ExecutorAgent involves a limited set of software or libraries. The code is then executed within the selected Docker image, providing isolated sandboxing that enhances security, reproducibility, and efficiency while eliminating the need for manual configuration overhead.

### Comparison with Other Bioinformatics Agents

To benchmark ToolsGenie 2.0’s performance against state-of-the-art bioinformatics agents, we conducted a comparative analysis with Biomni using two distinct benchmarks: an in-house curated dataset and the publicly available BixBench dataset (version 1.5)^15^. The in-house dataset comprises 140 open-ended bioinformatics data analysis tasks spanning key domains, including data processing, omics analysis, functional annotation, structural bioinformatics, and synthetic biology (Supplementary Table 1). BixBench consists of 205 questions drawn from real-world bioinformatics scenarios, aimed at evaluating agents’ proficiency in practical biological dataset exploration, data analysis, multi-step analytical workflows, and result interpretation (Supplementary Table 2)^15^. To ensure comparability, both ToolsGenie 2.0 and Biomni were configured with the Claude-4-Sonnet model. Each agent received identical queries and input files from the benchmarks, with no external modifications to their default behaviors. Average cost was calculated as the mean API usage cost per task under identical model and execution settings.

Performance on BixBench was measured by accuracy in its multiple-choice format, defined as the proportion of correctly selected options. For the in-house dataset, output correctness was independently assessed by three experienced bioinformaticians. To minimize biases from subjective scoring, evaluations were binary: 1 (for correct reasoning and a reasonable answer) or 0 (for incorrect or flawed outputs). For each erroneous response, the bioinformaticians provided brief descriptions of the underlying issues, as detailed in Supplementary Table 1.

## Acknowledgments

We thank the software engineering team at PromptBio for their support in deploying ToolsGenie to the cloud and integrating it into the PromptBio platform. We also thank our external trial users for their valuable feedback, which directly contributed to improving ToolsGenie.

